# BoNTax: a curated genome-based framework for taxonomic classification and characterization of *Clostridium botulinum* Group I and *Clostridium sporogenes*

**DOI:** 10.64898/2026.10.06.757027

**Authors:** Gangiredla Jayanthi, Shashi Sharma, Gonzalez-Escalona Narjol

## Abstract

*Clostridium botulinum* Group I and *Clostridium sporogenes* are closely related genomic lineages for which historical toxin-based nomenclature can obscure species-level relationships. We developed BoNTax, a curated taxonomy-first framework that separates average nucleotide identity (ANI)-based species assignment from botulinum neurotoxin (BoNT) characterization while integrating assembly quality assessment, multilocus sequence typing (MLST), AMRFinderPlus-based detection of *bont* genes, other virulence-associated genes, and antimicrobial resistance genes into standardized outputs. Analysis of 966 genomes informed genome-size criteria and the selection of 46 representative references comprising 22 *C. botulinum* Group I and 24 *C. sporogenes* genomes. Pairwise ANI resolved these references into two distinct species clusters: within-species ANI values were >95%, whereas between-species values were <95%, supporting a 95% species-level assignment threshold. BoNTax was implemented in GalaxyTrakr as complementary FASTQ- and FASTA-input workflows that converge on a common downstream analysis. The workflow completed analysis of all 42 validation genomes. Among the 32 genomes with a reported taxonomic identity, 29 assignments were concordant and three prompted taxonomic re-evaluations. For the ten genomes reported as taxonomically unknown, BoNTax provided ANI-based assignments. Of the three discordances, BT-22100019 was assigned to *C. sporogenes* rather than the reported *C. botulinum* Group I, whereas R1125/03 and R1135/03 showed 99.96% ANI to *C. caseinilyticum* DSM 46279, supporting their identification as *C. caseinilyticum*. MLST assignments were concordant for genomes with established profiles except PNUSAB000061, for which ST183 was not assigned by the MLST database version used in this analysis. BoNT type and subtype detection was fully concordant across toxigenic genomes, including bivalent strains. Thirty-one genomes met the predefined assembly-size QC criterion, whereas 11 received QC warnings, including all eight datasets representing four known mixed cultures. BoNTax therefore provides a reproducible genomic framework for resolving closely related clostridial lineages, identifying taxonomic annotations requiring re-evaluation, and reporting clinically relevant toxin information without conflating toxin genotype with species identity.

**Data Summary:** Accession numbers and associated metadata for the *Clostridium botulinum* Group I and *Clostridium sporogenes* genomes analyzed in this study are provided in Supplementary Table 1. This table includes the nucleotide, experiment, sample, and study accessions available for each genome. The complete pairwise average nucleotide identity matrix used to assess genomic diversity and establish the taxonomic decision criteria is provided in Supplementary Table 2. The distribution of curated *bont* profiles for the genomes in Supplementary Table 1 is provided in Supplementary Table 3. Integrated BoNTax results for the validation genomes and reference controls are provided in Supplementary Table 4 and include genome assembly metrics, quality-control status, antimicrobial resistance determinants, *bont* and other virulence-associated genes, multilocus sequence types, closest reference matches, ANI values, and taxonomic assignments.

The BoNTax FASTQ- and FASTA-input Galaxy workflow files, the versioned curated reference-genome manifest, a test dataset, expected outputs, software and database version information, and supporting documentation used in this study are archived in Zenodo as BoNTax version 1.0.0 (DOI: 10.5281/zenodo.22964346). The version-controlled development repository is maintained on GitHub (https://github.com/JayanthiGangiredla/BoNTax/tree/main). The step-by-step GalaxyTrakr protocol is available through Protocols.io (DOI: 10.17504/protocols.io.dm6gp4ozjgzp/v1). Sequence data used in this study are publicly available through the individual accession numbers listed in Supplementary Table 1.

**Impact Statement:** Historical toxin-based nomenclature can obscure the genomic boundary between *Clostridium botulinum* group I and *Clostridium sporogenes*, complicating taxonomic interpretation and surveillance reporting. BoNTax combines curated reference genomes and average nucleotide identity-based species assignment with assembly-size quality assessment, multilocus sequence typing, and detection of botulinum neurotoxin, other virulence-associated, and antimicrobial resistance genes. Complementary FASTQ- and FASTA-input GalaxyTrakr workflows make this integrated analysis accessible to laboratories working with either raw sequencing reads or assembled genomes. Analysis of a genomically diverse collection demonstrated clear separation of the two target taxa and identified public records requiring taxonomic re-evaluation. By independently reporting species identity and toxin genotype in standardized outputs, BoNTax supports reproducible interpretation of closely related clostridial genomes for food-safety, public-health, and research applications.

## 1. Introduction

Botulinum neurotoxin-producing clostridia represent a complex group of phylogenetically distinct organisms whose historical classification has been driven largely by toxin production rather than genomic relatedness [1]. Advances in comparative genomics have demonstrated that organisms currently designated as *C. botulinum* comprise several genetically divergent lineages that are more closely related to other *Clostridium* species than to one another [2, 3]. In particular, *C. botulinum* Group I and *C. sporogenes* exhibit substantial genomic similarity and overlapping biological characteristics, complicating species delineation when toxin carriage alone is used for classification [3, 4].

This has created an important distinction between clinical/public health nomenclature and microbial taxonomy. Clinical and surveillance frameworks, including those used in botulism investigations, often classify neurotoxin-producing isolates functionally as *C. botulinum* because of their pathogenic potential [5]. In contrast, modern microbial taxonomy increasingly supports classification based on core genome phylogeny and overall genomic relatedness, independent of mobile toxin gene content [3]. Because botulinum neurotoxin genes may be plasmid- or phage-associated and horizontally transferred, toxin production alone may not accurately reflect species-level evolutionary relationships [6–9].

Existing genomic tools can independently assess assembly quality, calculate genome similarity, identify sequence types, and detect toxin or resistance genes; however, a standardized framework that integrates these analyses while explicitly separating genome-based taxonomy from BoNT genotype is lacking for the closely related *C. botulinum* Group I and *C. sporogenes* lineages. To address the challenges associated with genomic characterization of botulinum neurotoxin-producing clostridia, we developed BoNTax, a curated genomic decision framework for the rapid identification and characterization of *C. botulinum* Group I and *C. sporogenes* from either raw sequencing reads or preassembled genomes (Figure 1).

**Figure 1.**
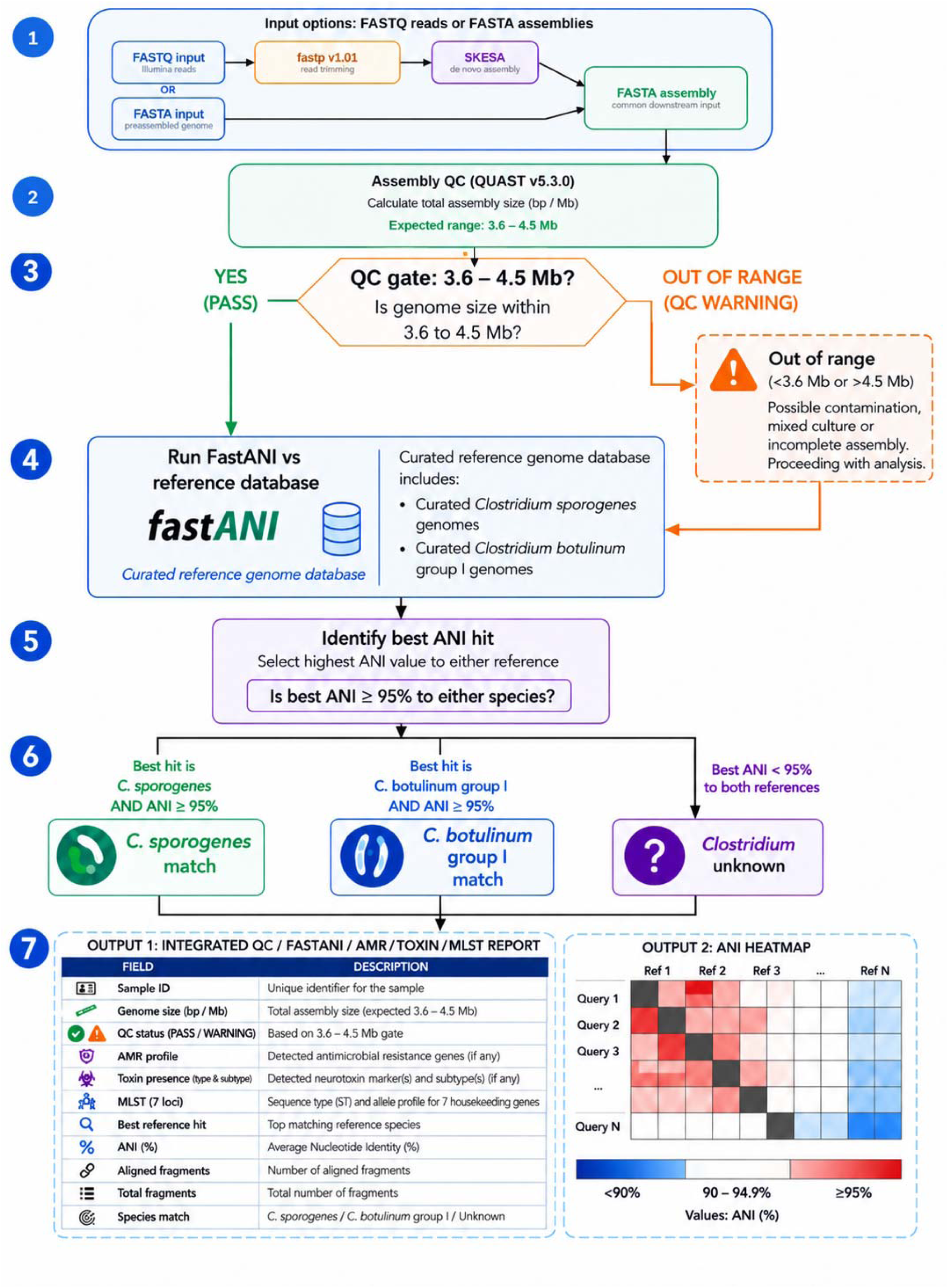
Curated genomic decision framework for rapid identification, toxin characterization, and quality assessment of *C. botulinum* Group I and *C. sporogenes*. The workflow integrates assembly QC, ANI-based taxonomic classification using curated reference genomes, toxin subtyping, AMR profiling, and automated reporting/visualization. Genome assemblies meeting the predefined size-based QC criterion (3.6–4.5 Mb) are compared against curated *C. botulinum* Group I and *C. sporogenes* reference genomes using FastANI, with species assignment based on a ≥95% ANI threshold. Genome assemblies outside the expected size range receive a QC warning but continue through the downstream ANI, MLST, and gene-detection modules; their results should be interpreted cautiously. Outputs include an integrated QC/ANI/MLST/BoNT/virulence/AMR report and an ANI heatmap.

BoNTax itself provides two dedicated entry points: a FASTQ workflow that processes raw sequencing reads through genome assembly and downstream characterization, and a FASTA workflow that analyzes preassembled genomes. Designed to improve standardization and reproducibility within genomic surveillance environments, both workflows incorporate assembly quality control, ANI-based taxonomic classification against curated reference genomes, toxin gene detection and subtyping, antimicrobial resistance profiling, and automated reporting and visualization. Importantly, BoNTax prioritizes genomic taxonomy for species assignment while independently reporting neurotoxin gene content and subtype information, thereby separating taxonomic classification from clinically relevant toxin characterization.

## 2. Materials and methods

### 2.1 BoNTax workflow architecture and implementation

A high-level overview of BoNTax is presented in Figure 1. The BoNTax workflows are accessible through the FDA HFP GalaxyTrakr graphical user interface (https://galaxytrakr.org) [10, 11]. A step-by-step protocol for executing the BoNTax_FASTA and BoNTax_FASTQ workflows is available through Protocols.io (https://dx.doi.org/10.17504/protocols.io.dm6gp4ozjgzp/v1). The BoNTax FASTQ- and FASTA-input Galaxy workflow files, versioned curated reference-genome manifest, test dataset, expected outputs, software and database version information, and supporting documentation used in this study are archived in Zenodo as BoNTax version 1.0.0 (DOI: 10.5281/zenodo.22964346). The BoNTax FASTQ- and FASTA-input Galaxy workflow files, the versioned curated reference-genome manifest, a test dataset, and documentation required to reproduce the analyses are available at GitHub (https://github.com/JayanthiGangiredla/BoNTax/tree/main).

The BoNTax pipeline accommodates either raw sequencing reads (FASTQ) or preassembled genomes (FASTA). The FASTQ workflow trims reads using *fastp* v1.01 [12] and assembles them with SKESA v2.3.0 [13] before converging with the FASTA workflow. Both input pathways utilize identical downstream modules: QUAST v5.3.0 [14] for assembly quality control (QC); FastANI v1.3 [15] for average nucleotide identity (ANI)-based taxonomic classification against a curated reference collection; mlst v2.22.0 [16] for multilocus sequence typing [17]; and AMRFinderPlus v4.2.5 [18] for detecting *bont* genes, other virulence-associated factors, and antimicrobial resistance (AMR) determinants.

Module outputs are automatically parsed into two principal files: (1) a consolidated QC_ANI_results table summarizing QC status, ANI-based taxonomy, sequence type, and virulence/AMR profiles; and (2) a PDF heatmap visualizing the ANI-based clustering of query genomes relative to the *C. botulinum* Group I and *C. sporogenes* reference groups. This modular architecture eliminates the need for command-line execution, standardizes reporting, and allows individual databases to be easily updated as genomic resources evolve.

### 2.2 Curated reference genome selection and ANI classification criteria

A curated reference genome dataset comprising twenty-two *C. botulinum* Group I and twenty-four *C. sporogenes* assemblies (Table 1) was constructed from publicly available NCBI genomes and selected from a larger collection of 966 *C. botulinum* Group I and *C. sporogenes* genomes (Supplementary Table 1). Genomes were selected based on assembly quality, taxonomic consistency, genomic representation, and availability of toxin characterization metadata. Reference genomes were also selected to represent the observed intra-species ANI diversity range (95–100%) (Supplementary Table 2). These reference genomes were used to anchor ANI-based taxonomic assignment and to evaluate genomic separation between the two taxa. For each query assembly, the highest ANI value across all curated reference genomes was retained. An ANI threshold of ≥ 95% was used for species-level assignments [15, 19, 20]. A genome query was assigned to the species represented by its best-matching reference when ANI was ≥95%. Genome queries with a best ANI value <95% were reported as *Clostridium* sp. unknown. The query-by-reference ANI matrix was generated by pivoting FastANI output, with mean aggregation applied only to duplicate query–reference records. Reciprocal ANI values were retained as separate matrix entries. Rows and columns of the untransformed matrix were clustered using Euclidean distances between ANI-profile vectors and average-linkage clustering in Galaxy heatmap2 v3.3.0+galaxy0. The resulting dendrograms represent clustering of ANI profiles and not formal phylogenetic relationships.

**Table 1.** Representative *C. botulinum* Group I and *C. sporogenes* strains selected to span the observed intra-species ANI diversity range (95–100%), with ANI values calculated relative to the representative genomes Hall 183 (*C. botulinum* Group I) and ATCC 15579 (*C. sporogenes*).

| *C. botulinum* |  |  |
| --- | --- | --- |
|  | Group I | *C. sporogenes* |
| Representatives *C. botulinum* | ANI (%) vs. Hall 183 | ANI (%) vs ATCC 15579 |
| Group I |  |  |
| Colworth BL30 | 95.98 | 92.57 |
| A2B7 92 | 96.12 | 92.64 |
| PNUSAB000059 | 96.79 | 93.02 |
| CDC64197 | 97.31 | 92.96 |
| CDC14 | 98.23 | 92.93 |
| A96 | 98.42 | 93.01 |
| CDC43305 | 99.11 | 92.92 |
| PNUSAB000004 | 99.42 | 93.00 |
| CDC28023 | 99.8 | 92.90 |
| CDC67190 | 100 | 93.14 |
| 657 | 96.65 | 93.13 |
| Bf | 96.66 | 93.09 |
| Hall 183 | 100 | 93.04 |
| Hall 4834 | 99.92 | 92.94 |
| Hall 80 | 97.58 | 92.99 |
| Hall 8388A | 99.99 | 93.05 |
| Hall 8857Ab | 99.91 | 93.00 |
| Hall | 97.53 | 93.11 |
| Kyoto | 98.44 | 93.05 |
| Langeland | 97.27 | 93.01 |
| Loch Maree A3 | 96.07 | 92.71 |
| NCTC 2916 | 99.99 | 93.12 |
| **Representatives C.** | **ANI (%) vs. Hall 183** | **ANI (%) vs ATCC** |
| ***sporogenes*** |  | **15579** |
| 2015006229-DFPST | 93.25 | 96.71 |
| 87-0535 | 92.09 | 95.11 |
| 88-0163 | 92.93 | 99.30 |
| AM1194 | 92.84 | 96.95 |
| AM553 | 93.10 | 99.36 |
| ATCC 15579 | 93.08 | 100.00 |
| B2 450 | 93.10 | 99.32 |
| CDC38839 | 92.82 | 96.96 |
| CDC45459 | 92.77 | 96.94 |
| CDC67240 | 93.23 | 96.84 |
| CDC75088T1 | 93.28 | 97.02 |
| CDC76054PL1 | 93.13 | 97.35 |
| CDC76098 | 93.23 | 96.93 |
| CDC78112PL1 | 93.27 | 96.92 |
| CDC_1632 | 93.46 | 96.15 |
| CDC_31846 | 93.16 | 97.42 |
| DSM 795 | 92.09 | 95.12 |
| FDA1304820-S005-002 | 92.25 | 95.34 |
| FDA1316851-S005-003 | 92.03 | 95.23 |
| FDA1316851-S005-004 | 92.06 | 95.14 |
| Osaka05 | 92.81 | 96.99 |
| PA 3679 | 92.09 | 95.01 |
| Prevot_594 | 92.09 | 95.20 |
| W534242 | 93.20 | 96.73 |

### 2.3 Assembly quality-control criteria

Assembly QC thresholds were established based on expected genome size distributions. Assemblies with genome sizes below 3.6 Mb or above 4.5 Mb were flagged as potential mixed cultures, contamination, or incomplete assemblies.

### 2.4 Genome-wide characterization of *bont* profiles

Genome assemblies (Supplementary Table 1) were retrieved from GenBank. Initial *bont* gene annotations were sourced from NCBI Pathogen Detection (https://www.ncbi.nlm.nih.gov/pathogens/ accessed July 1^st^ 2026) [18, 21]. For botulinum neurotoxin and antimicrobial resistance (AMR) gene detection, when multiple hits represented the same determinant, the highest-confidence hit was retained; distinct qualifying genes, including multiple *bont* genes in bivalent genomes, were reported individually. Assignments were refined using MBioSEQ Ridom Typer v12.0.5 (Bruker) (https://www.ridom.de/ridom-typer/), against a curated reference set (requiring ≥90% nucleotide identity and ≥99% coverage) to resolve ambiguous, partial, or type-only NCBI calls [22]. Candidate sequences were retained when they showed ≥90% nucleotide identity across ≥99% of the reference length. Among retained candidates, subtype assignment was based on the highest-scoring full-length reference match and was manually reviewed when the NCBI Pathogen Detection and in-house assignments differed. Final curated profiles (e.g., A1, or A1B5 for multiple genes) reconciled both methods. This independent subtype-curation procedure was used to characterize the 966-genome source collection and was separate from the automated BoNTax Galaxy workflows, in which *bont* genes are detected and reported by AMRFinderPlus. Assemblies lacking *bont* genes were designated “No toxin gene detected” (ND), noting that phenotypic toxicity was not evaluated. Results are detailed in Supplementary Table 1 and summarized by ANI-defined taxonomic group in Supplementary Table 3.

### 2.5 Validation Genome Dataset

A validation dataset consisting of forty-two genomes was compiled from NCBI, FDA isolate collections, and additional external datasets, with some genomes carried forward from the reference dataset (Table 2). The validation set included toxigenic *C. sporogenes*, non-toxigenic *C. sporogenes*, *C. botulinum, Clostridium* species other than *C. botulinum* or *C. sporogenes*, isolates of unknown taxonomic identity, and mixed-culture assemblies. Represented toxin subtypes included A1, bivalent A1b5 (lower case indicate that this toxin gene is silent or inactive due to a mistranslation), A2, A3, B1, B5, bivalent B5F2, F3, F5, and X. This validation dataset was designed to assess workflow robustness, real-world applicability, and performance on biologically challenging isolates. Validation genomes were used to evaluate workflow performance, classification accuracy, edge-case handling, and reproducibility.

**Table 2.**
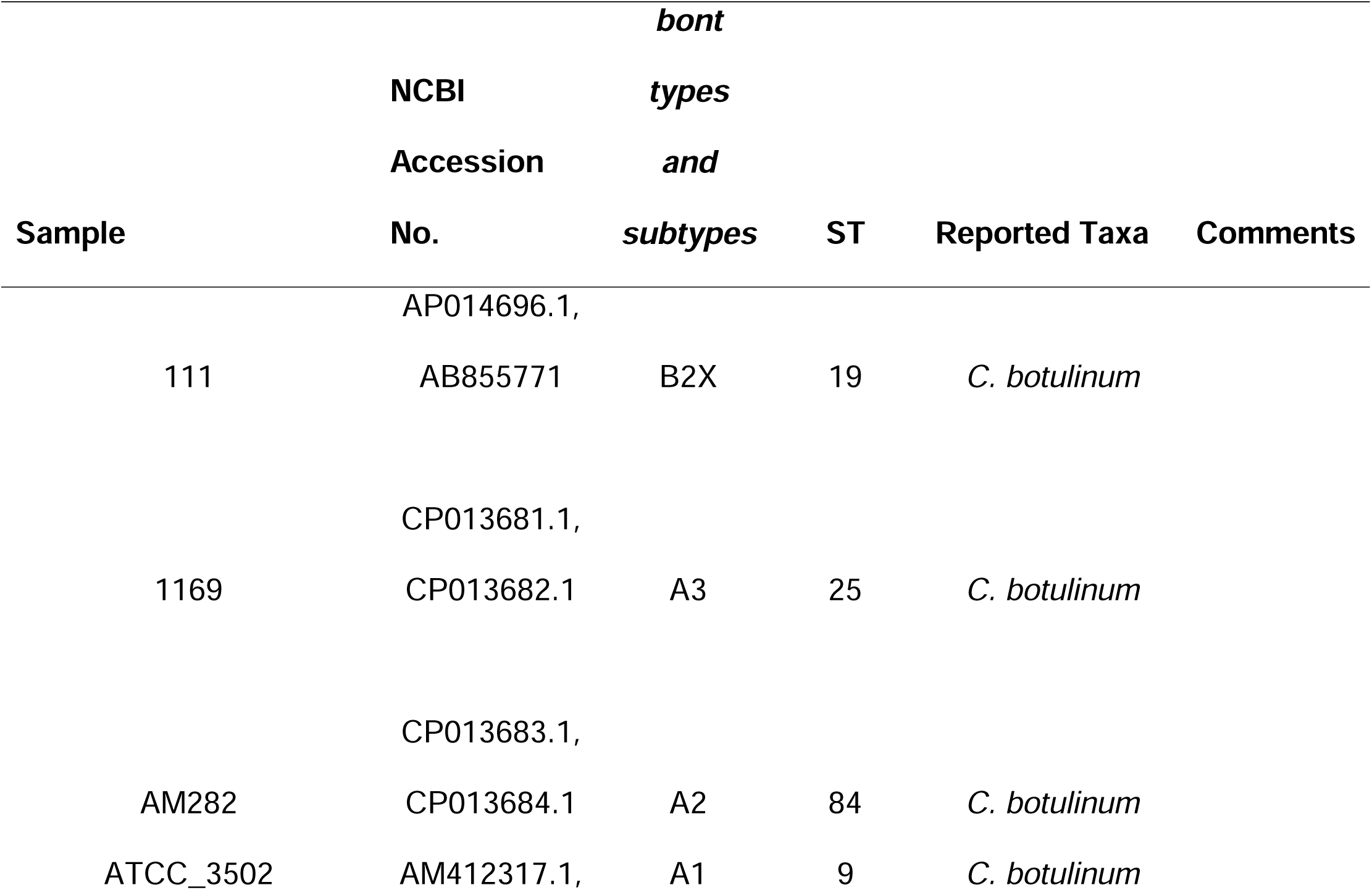
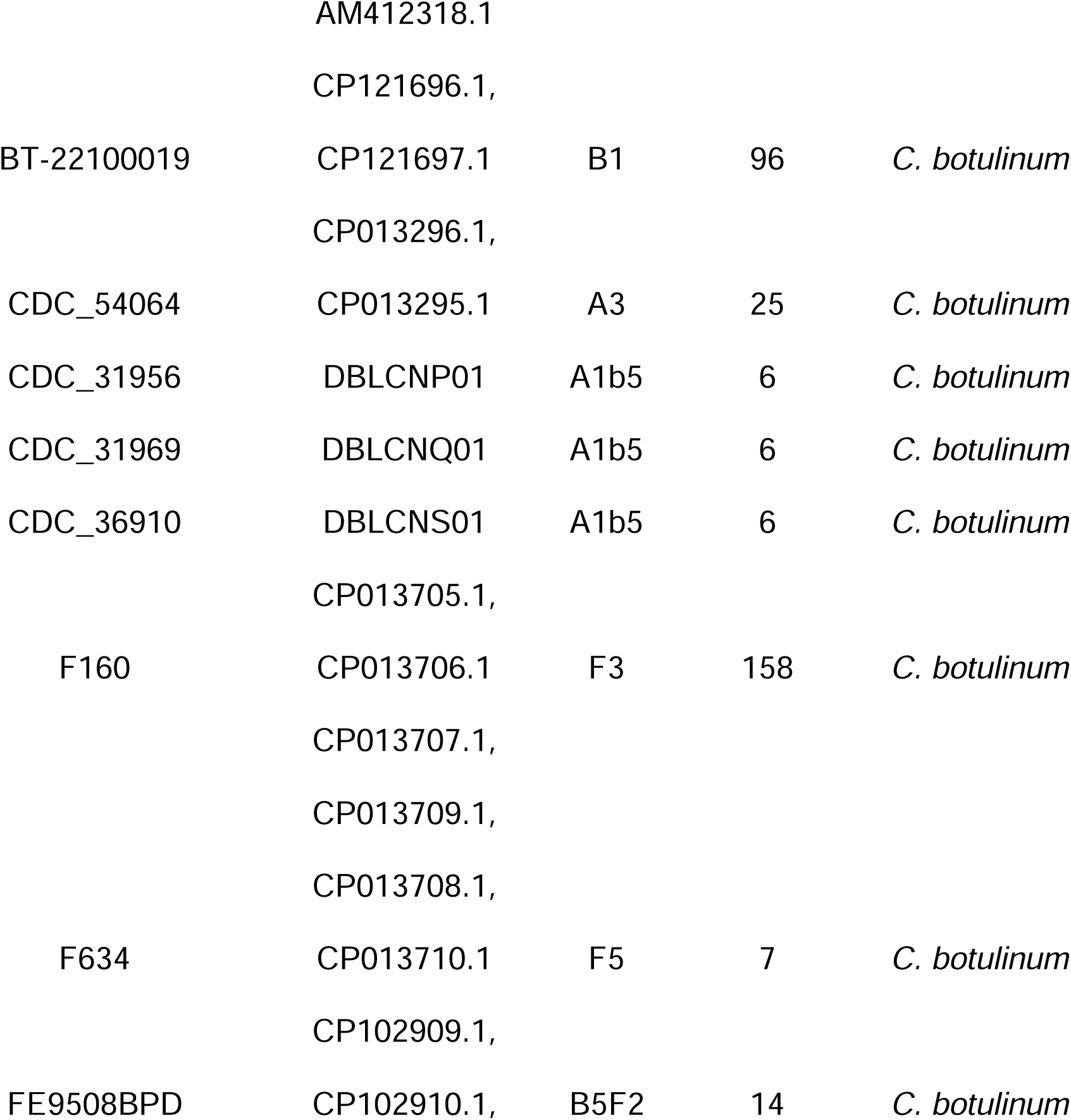
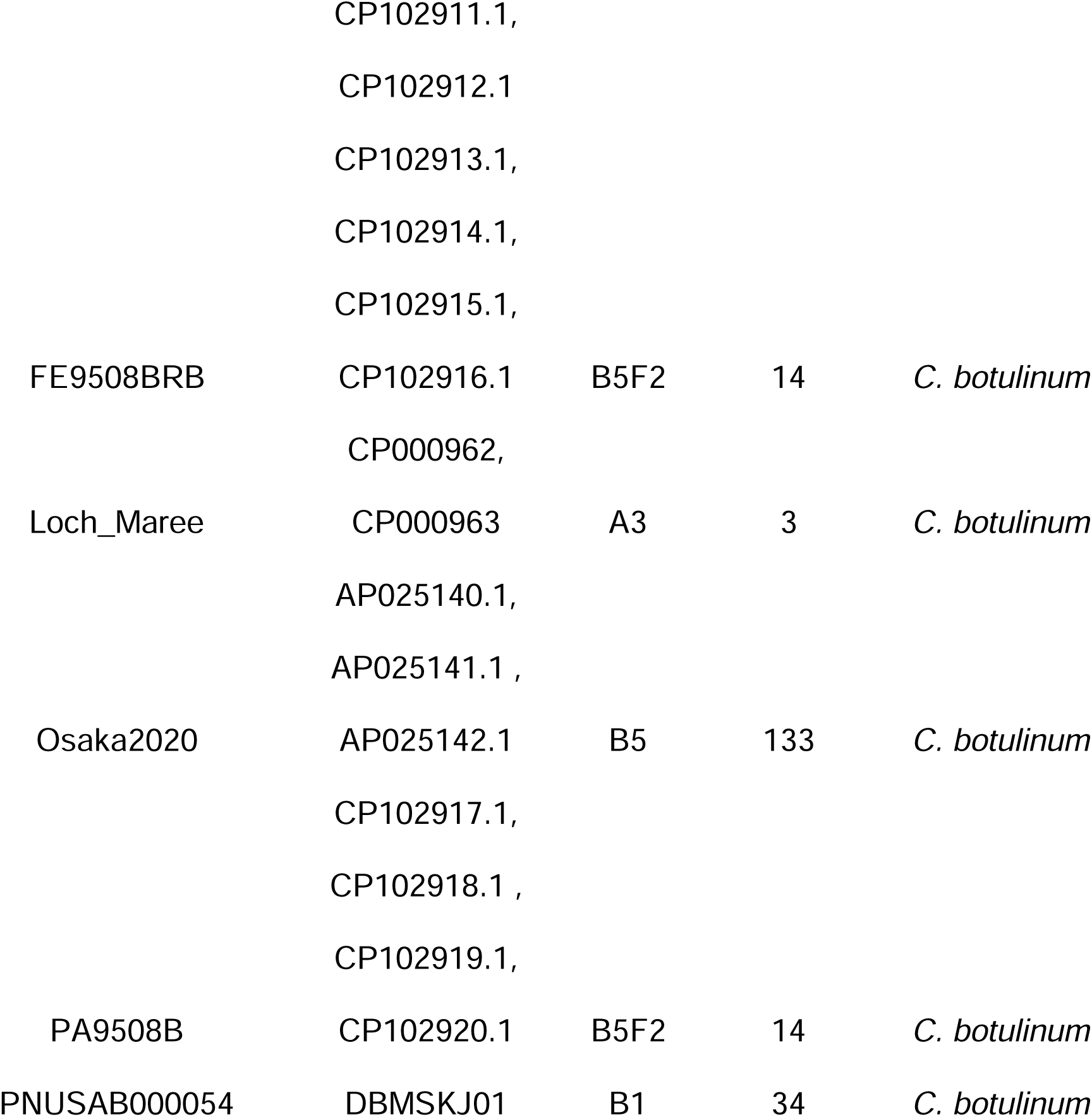
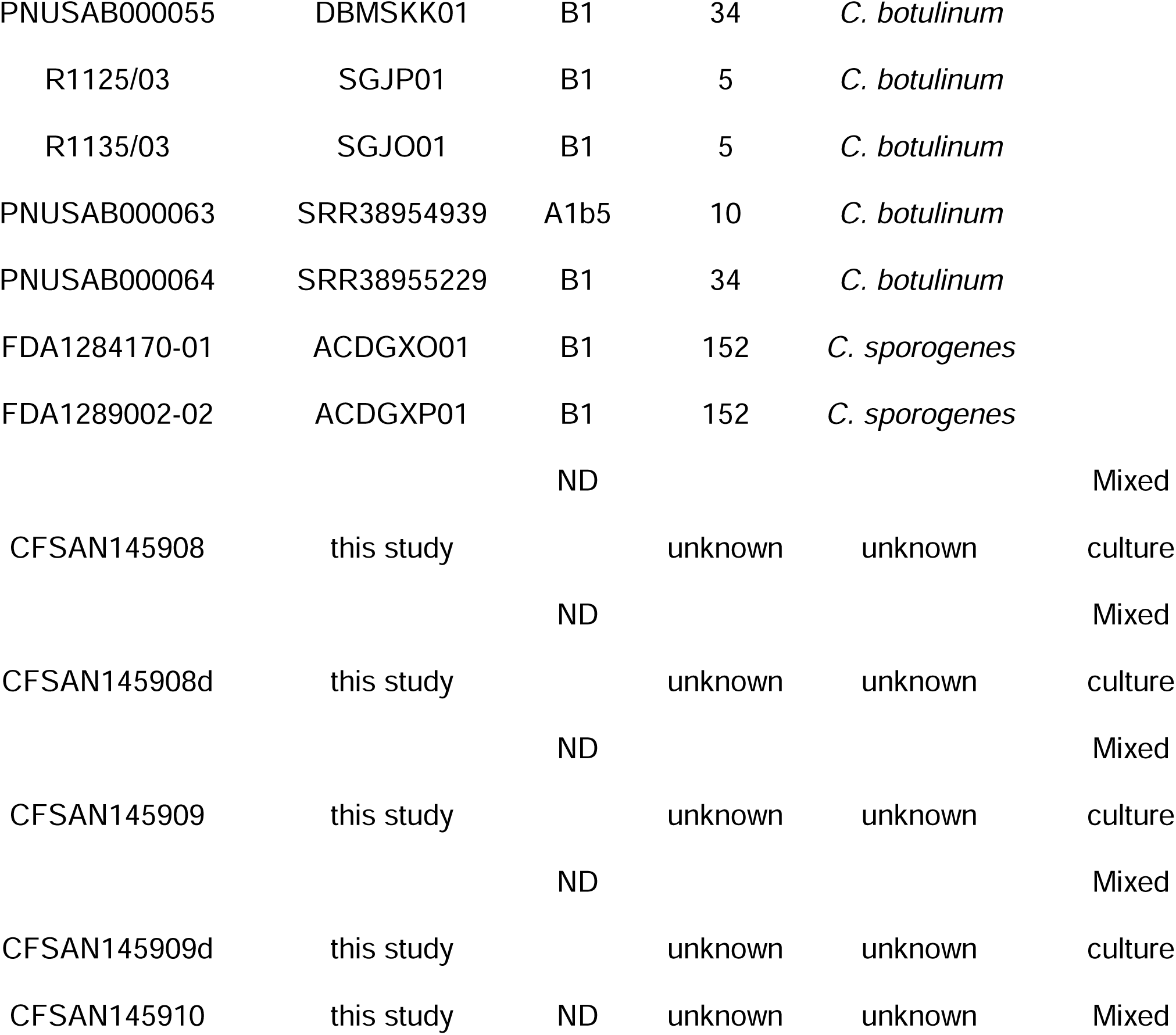
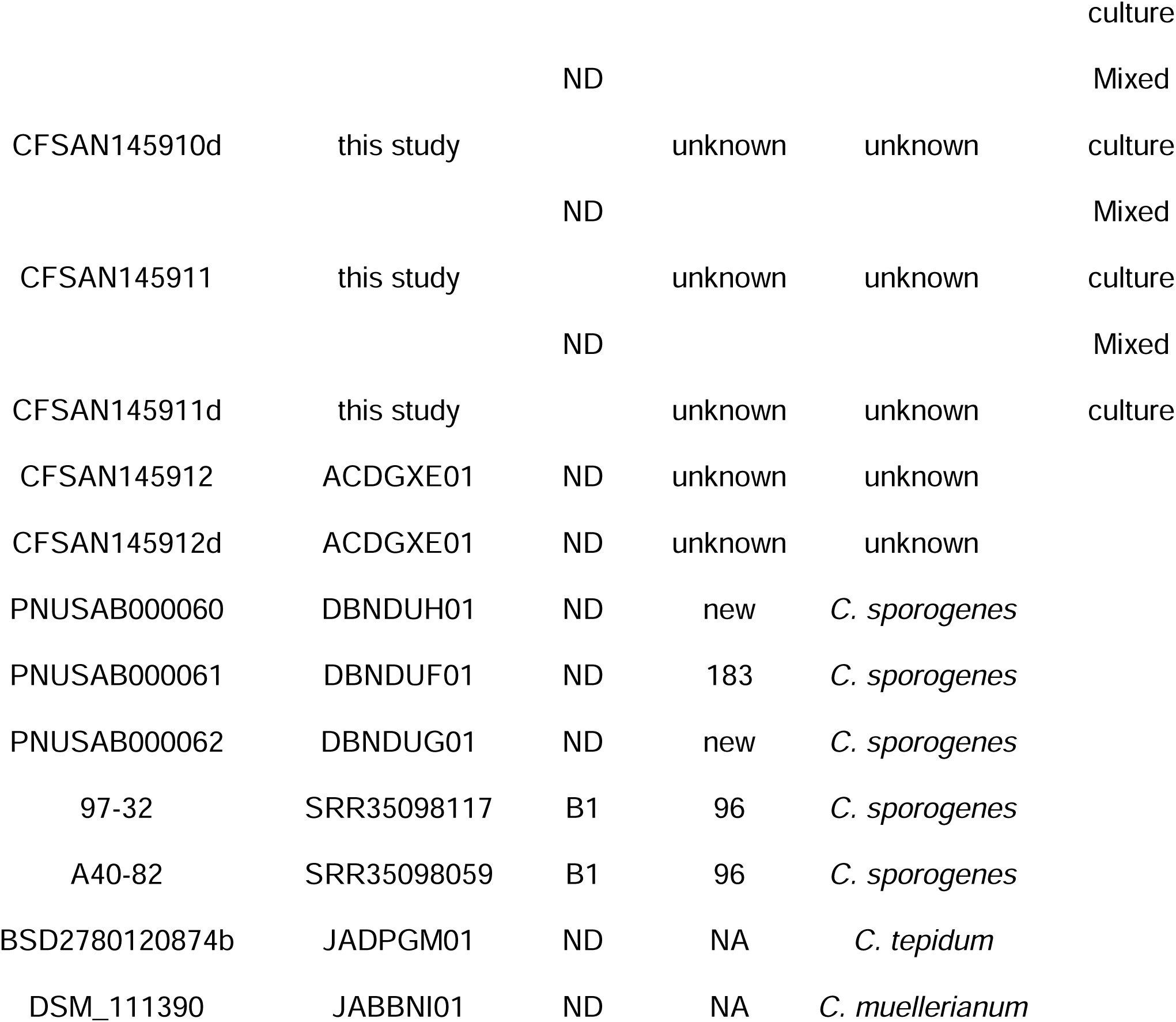
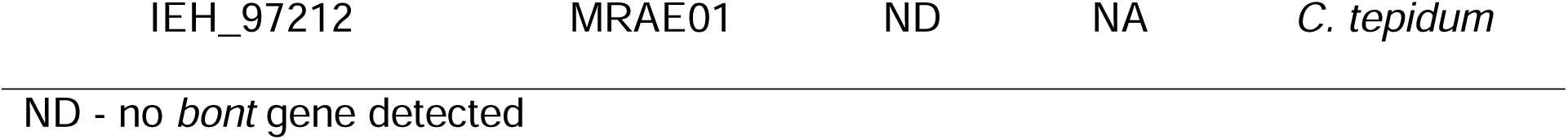
Forty-two validation genomes used to validate the automated classification and reporting functions of the BoNTax pipeline. The validation set consisted primarily of *C. botulinum* and *C. sporogenes* genomes and included additional *Clostridium* species to evaluate taxonomic discrimination.

## 3. Results

### 3.1 Genome-size distributions and ANI support assembly screening and species-level separation of *C. botulinum* Group I and *C. sporogenes*

Total assembly sizes across the curated 966-genome collection were concentrated between 3.6 and 4.5 Mb; this empirically observed interval was adopted as the assembly-size screening criterion (Supplementary Table 1), supporting the implementation of assembly-size quality control thresholds. ANI comparisons demonstrated a clear separation between *C. botulinum* Group I and *C. sporogenes*, supporting the use of a 95% ANI threshold for species assignment (Supplementary Table 2).

### 3.2 Curated reference genomes resolve the genomic boundary between *C. botulinum* Group I and *C. sporogenes*

The selected genomes for the reference dataset encompassed the observed intra-species genomic diversity of both species, spanning ANI values from approximately 95% to 100% relative to the selected representative genomes (Table 1). The dataset included genomically diverse isolates representing major lineages within *C. botulinum* Group I and *C. sporogenes*. Pairwise ANI analysis of the selected reference genomes revealed a clear genomic separation between *C. botulinum* Group I and *C. sporogenes* (Figure 2). Hierarchical clustering of the ANI values resolved the genomes into two distinct species-associated clusters. All within-species comparisons had ANI values ≥95%, whereas all comparisons between the two species had ANI values <95%. This non-overlapping separation supports the taxonomic coherence of the curated reference groups and the use of a ≥95% ANI threshold for species assignment in BoNTax. The resulting dendrograms represent similarity-based clustering of ANI profiles rather than formal phylogenetic relationships.

**Figure 2.**
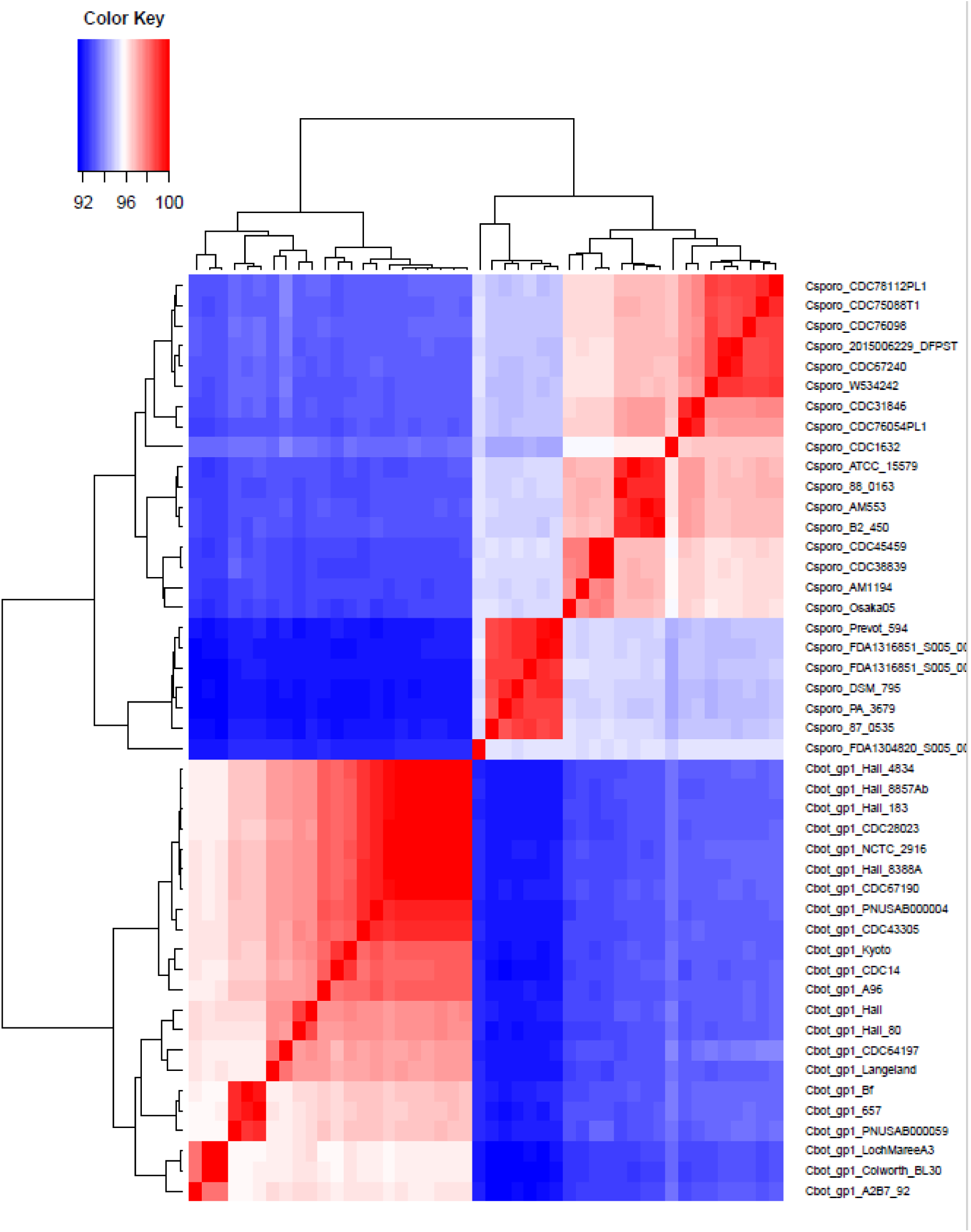
ANI-based genomic delineation of *C. botulinum* Group I and *C. sporogenes* reference genomes. Heatmap of average nucleotide identity (ANI) values among curated reference genomes representing the diversity of both species. The resulting clustering resolves the genomes into two distinct species groups. Within-species ANI values (≥95%) clearly contrast with between-species values (<95%), supporting the ≥95% ANI threshold for taxonomic assignment in BoNTax. Dendrograms represent similarity-based clustering of ANI profiles, not formal phylogenetic relationships.

### 3.3 Distribution of *bont* gene profiles across ANI-defined taxa

Across the 966-genome collection, 938 genomes were assigned by ANI to *C. botulinum* Group I and 28 to *C. sporogenes*. A *bont* gene was identified in 912 *C. botulinum* Group I genomes and twenty *C. sporogenes* genomes, whereas 26 and 8 genomes, respectively, had no *bont* gene detected (Supplementary Table 3). The *C. botulinum* Group I genomes contained a broad diversity of A-, B-, and F-associated profiles, including genomes harboring more than one *bont* gene. Notably, the B1, B2, B3, B5, and B6 profiles occurred in both ANI-defined taxa (Supplementary Table 3). Thus, genomes with the same BoNT subtype could be assigned to either *C. botulinum* Group I or *C. sporogenes*, indicating that toxin genotype did not uniquely correspond to genome-based species identity.

### 3.4 Validation of automated classification and reporting

To validate the BoNTax pipeline, we selected 42 validation genomes. Most of them belonged to either *C. botulinum* or *C. sporogenes* species. These genomes have been deposited at GenBank (Table 2), except for samples CFSAN145908, CFSAN145909, CFSAN145910, and CFSAN145911 which were mixed culture samples. These datasets were included to evaluate the performance and potential utility of BoNTax for the analysis of mixed cultures. The *C. botulinum* or *C. sporogenes* genomes were highly diverse and encompassed numerous genomically distinct groups and diverse botulinum neurotoxin gene content, thereby increasing the complexity of the validation dataset.

The BoNTax pipeline successfully analyzed all 42 validation genomes. In addition, eight reference genomes included as controls for pipeline execution and visualization (*C. botulinum* Group I: CDC67190, Hall_80, Langeland, and Loch Maree A3; *C. sporogenes*: ATCC 15579, DSM 795, Osaka05, and Prevot 594) were correctly classified (Supplementary Table 4 and Supplementary Figure 1). Comparison of the BoNTax output with the expected reference metadata demonstrated a high level of concordance for species assignment, sequence type (ST), and botulinum neurotoxin (BoNT) gene content. Among the 32 validation genomes with a reported taxonomic identity, 29 BoNTax assignments were concordant and three prompted taxonomic re-evaluations. Three genomes showed discrepancies with the reference metadata. BT-22100019 was identified as *C. sporogenes* by BoNTax, matching a *C. sporogenes* strain CDC31846 at 99.92% ANI, whereas it was reported as *C. botulinum* Group I in the reference dataset [23]. In addition, R1125/03 and R1135/03, which were also reported as *C. botulinum* in the reference metadata, were not classified within the *C. botulinum* Group I or *C. sporogenes* reference groups by BoNTax. Subsequent ANI analysis showed that both genomes matched *Clostridium caseinilyticum* strain DSM 46279 [24] at 99.96% ANI, supporting their assignment to *C. caseinilyticum* rather than *C. botulinum*. These results support taxonomic re-evaluation of the three reported *C. botulinum* annotations. For the remaining ten validation genomes reported as taxonomically unknown, BoNTax provided ANI-based assignments.

ST assignments were concordant for genomes with previously defined MLST profiles, except PNUSAB000061, for which the expected ST183 profile was not assigned by the version of the MLST database used in this analysis. For several genomes annotated as having an unknown ST in the reference dataset, BoNTax assigned sequence types, including CFSAN145908 (ST72), CFSAN145911 (ST70), and CFSAN145912 (ST74) and their respective duplicate datasets. Conversely, genomes annotated as possessing novel sequence types ("new") were not assigned an ST by BoNTax, as expected, because these sequence types are not represented in the current *C. botulinum* MLST database (https://pubmlst.org/organisms/clostridium-botulinum).

BoNT gene detection was fully concordant with the expected toxin profiles, with BoNTax correctly identifying the BoNT toxin types and subtypes for all toxigenic genomes. This included genomes harboring more than one BoNT gene, for which the corresponding toxin types and subtypes were correctly identified. Although BoNTax reports individual *bont* genes and their subtypes (e.g., *bont/A1* and *bont/B5*) rather than the condensed notation used in the reference metadata (e.g., A1B5), the toxin type and subtype assignments were concordant. For non-toxigenic genomes, BoNTax correctly reported the absence of *bont* genes and, when present, identified other clostridial virulence-associated genes, including *cpa*, *cpb2*, and *pfoA*.

Assembly-size quality control indicated that 31 of the 42 validation genomes met the predefined assembly-size QC criterion, whereas 11 were flagged as “Possible contamination or Mixed culture or an incomplete assembly”. The flagged datasets included eight assemblies representing four known mixed cultures (CFSAN145908, CFSAN145909, CFSAN145910, and CFSAN145911 and their respective duplicates) and three genomes belonging to species other than *C. botulinum* or *C. sporogenes* (strains BSD2780120874b, DSM_111390, and IEH_97212).

The ANI-based heatmap complemented the taxonomic assignments reported in the QC_ANI_results table by providing a visual representation of the relationships among the validation and reference genomes (Supplementary Figure 1). Genomes assigned to *C. botulinum* Group I or *C. sporogenes* clustered with their corresponding reference genomes, whereas genomes belonging to other *Clostridium* taxa were positioned outside these groups. The heatmap therefore provided visual support for the ANI-based taxonomic assignments and allowed genomes that did not belong to either *C. botulinum* Group I or *C. sporogenes* to be readily distinguished.

Overall, BoNTax showed high concordance with the expected taxonomic, MLST, and BoNT profiles across this genomically diverse validation dataset. The integrated assembly-size QC assessment and ANI-based visualization provided complementary information for flagging datasets consistent with mixed cultures, potential assembly-quality issues, and genomes outside the *C. botulinum* group I and *C. sporogenes* reference groups. Because the assembly-size QC interval was calibrated specifically for *C. botulinum* group I and *C. sporogenes*, valid genomes belonging to other *Clostridium* species may receive a warning. Such warnings should be interpreted together with the ANI-based taxonomic assignment rather than as definitive evidence of contamination or incomplete assembly. Moreover, discrepancies between pipeline assignments and reported metadata prompted further genomic investigation, as illustrated by R1125/03 and R1135/03, for which subsequent ANI analysis supported their identification as *C. caseinilyticum* rather than the previously reported *C. botulinum*.

## DISCUSSION

### A taxonomy-first framework for biologically complex *Clostridium* samples

This study describes a curated genomic decision framework for the rapid identification and characterization of *C. botulinum* Group I and *C. sporogenes*. Unlike traditional workflows that emphasize toxin production as the primary classification criterion [5], this framework prioritizes genome-based taxonomic assignment using curated reference genomes and ANI-based classification while independently reporting toxin and AMR profiles. The framework accommodates both raw sequencing reads and assembled genomes through dedicated FASTQ- and FASTA-input workflows, respectively, which converge on the same downstream taxonomic and genotyping analyses within GalaxyTrakr [10]. This separation of genome-based species identity from toxin genotype is central to BoNTax and addresses a longstanding challenge in the classification of BoNT-producing clostridia.

Validation of BoNTax across a genomically diverse collection of *Clostridium* genomes demonstrated the utility of integrating assembly quality, ANI-based taxonomy, genotyping, and toxin characterization within a single workflow. The combination of assembly-level QC and ANI-based classification allows taxonomic assignments to be interpreted in the context of genome quality, while the curated reference collection provides a consistent framework for distinguishing *C. botulinum* Group I from *C. sporogenes* and other *Clostridium* taxa. The identification of known mixed-culture datasets by the QC component further illustrates the importance of evaluating genotyping results together with indicators of assembly and sample quality. Importantly, the workflow determines taxonomic identity independently of BoNT gene content, allowing toxigenic *C. sporogenes* isolates to retain their genome-based species assignment while their BoNT type and subtype are reported separately.

### Clinical and Genomic Classification Frameworks

Clinical and public health investigations often prioritize toxin production because it directly relates to disease risk, outbreak response, and food safety decision-making [5]. Consequently, neurotoxin-producing isolates have historically been referred to operationally as *C. botulinum* based primarily on their ability to produce botulinum neurotoxin [5]. This approach reflects a disease-centered and toxin-centered framework in which the presence of a botulinum neurotoxin gene is the defining characteristic of interest. In contrast, the objective of the present framework is taxonomic classification based on genome-wide similarity and evolutionary relationships [2–4, 25]. This distinction is particularly relevant because *bont* genes occur in different genomic backgrounds, may be located on mobile genetic elements, and can undergo horizontal transfer [7–9]. Under this framework, an isolate whose genome-wide ANI places it within *C. sporogenes* should be classified as *C. sporogenes*, even if it carries a botulinum neurotoxin gene [3, 4, 26]. The toxin genotype is then reported as an independent characteristic (e.g., *C. sporogenes* BoNT/B1) rather than being used to redefine species identity. We argue that this approach more accurately reflects evolutionary relationships while preserving the toxin information necessary for clinical, food safety, and public health applications. Thus, the clinical and genomic frameworks should be viewed as complementary but distinct systems: one focused on pathogenic potential and disease risk, and the other focused on biological classification and evolutionary history.

This distinction was also evident across the broader 966-genome collection. BoNT/B subtypes B1, B2, B3, B5, and B6 were detected in both ANI-defined taxa (Supplementary Table 3), showing that BoNT genotype alone did not uniquely predict whether a genome belonged to *C. botulinum* Group I or *C. sporogenes*. This overlap provides empirical support for the independent reporting of ANI-based taxonomy and toxin genotype in BoNTax. Because the genome collection was assembled from available sequence datasets rather than through population-based sampling, the observed toxin-profile frequencies describe the composition of the analyzed collection and should not be interpreted as prevalence estimates for either species.

The validation results provide practical examples of the importance of separating historical organism designation from genome-based taxonomy. R1125/03 and R1135/03 were reported as *C. botulinum* in GenBank (BioSample SAMN10822427 and SAMN10822428) but were positioned outside the *C. botulinum* Group I and *C. sporogenes* reference groups by BoNTax. Subsequent ANI analysis identified *C. caseinilyticum* DSM 46279 [24] as the closest match for both genomes (99.96% ANI), supporting their classification as *C. caseinilyticum* rather than *C. botulinum*. Similarly, BT-22100019 was assigned to *C. sporogenes* by BoNTax despite being reported as *C. botulinum* Group I in GenBank [23]. These cases illustrate how genome-based classification can identify taxonomic annotations that warrant re-evaluation and reinforce the importance of reporting taxonomic identity and toxin genotype as related but distinct characteristics [3, 4].

### Utility of curated reference genomes and standardized decision criteria

Curated reference genomes are central to the performance and reliability of the BoNTax framework. Species assignment is influenced not only by ANI values but also by the taxonomic integrity and genomic breadth of the reference dataset. Public genome repositories contain records generated across different periods and under evolving taxonomic frameworks, and annotations may therefore not always reflect current genome-based relationships [3, 27]. Careful reference selection is consequently essential for robust interpretation [28]. By incorporating genomically representative genomes spanning the known diversity of *C. botulinum* Group I and *C. sporogenes*, the framework establishes a stable and reproducible basis for taxonomic classification. The application of standardized ANI thresholds further improves the consistency, transparency, and interpretability of classification decisions across laboratories and surveillance programs [15].

The integration of these components within GalaxyTrakr provides additional practical advantages for genomic surveillance. BoNTax consolidates assembly QC, taxonomic classification, MLST, BoNT characterization, AMR and other virulence-associated gene detection, and standardized reporting into an automated workflow. Its two input options allow laboratories to analyze either raw sequencing reads or existing genome assemblies while retaining a common downstream analytical framework. The modular Galaxy implementation also allows individual analytical tools, databases, and reference collections to be updated as genomic resources and taxonomic knowledge evolve. Together, these features can facilitate standardized interpretation and reporting across food safety, public health, and research laboratories.

Several limitations should nevertheless be considered. Taxonomic assignment remains dependent on the completeness, quality, and taxonomic accuracy of the reference genome collection and will require periodic reassessment as additional genomes and newly described taxa become available. Although ANI provides a practical and widely used measure of genome relatedness [15], isolates near species boundaries or outside the diversity represented by the curated reference collection may require additional phylogenomic investigation. Similarly, QC warnings generated by the workflow indicate potential contamination, mixed culture, or incomplete assembly but do not distinguish definitively among these conditions and should therefore prompt further evaluation rather than be interpreted as a specific diagnosis. Finally, because *bont* genes can occur in different genomic backgrounds, toxin content should not be considered definitive evidence of species identity [2, 7–9]. Future development of BoNTax will focus on expanding the curated reference collection to additional botulinum groups and newly described taxa, incorporating complementary phylogenomic approaches for ambiguous classifications, and extending the framework to support broader clostridial genomic surveillance.

## Conclusions

By integrating curated reference genomes and ANI-based species assignment with independent BoNT characterization, BoNTax provides a reproducible framework for distinguishing closely related *C. botulinum* Group I and *C. sporogenes* lineages without conflating taxonomic identity with toxin-associated pathogenic potential.

## Ethical approval and consent to participate

Not applicable. This study involved the analysis of bacterial genome assemblies and associated non-identifiable metadata and did not involve human participants, identifiable private human information, or live vertebrate animals. Institutional ethical approval and informed consent were therefore not required.

## Disclaimer

The views expressed in this article are those of the authors and do not necessarily reflect the official policy of the U.S. Food and Drug Administration. Reference to any commercial product, process, service, manufacturer, or company does not constitute or imply endorsement by the U.S. Government.

## Author contributions

**Jayanthi Gangiredla:** Methodology, Software, Data curation, Formal analysis, Investigation, Validation, Visualization, Writing – review and editing.

**Shashi Sharma:** Conceptualization, Methodology, Investigation, Writing – review and editing.

**Narjol Gonzalez-Escalona:** Conceptualization, Methodology, Data curation, Formal analysis, Investigation, Resources, Supervision, Project administration, Validation, Visualization, Writing – original draft, Writing – review and editing.

All authors reviewed and approved the final version of the manuscript and agreed to be accountable for the work.

## Acknowledgements

The authors thank the FDA Human Foods Program for providing resources for the investigation. OpenAI ChatGPT (GPT-5.6 Sol Pro; used August–September 2026) was used to assist with manuscript organization, language editing, and consistency checking. The authors directed its use, independently verified the scientific content and interpretation, and reviewed and approved the final manuscript. The tool was not used to generate primary sequence data or to make final scientific conclusions.

## SUPPLEMENTARY MATERIAL LEGENDS

Supplementary Figure 1. ANI-based placement of the 42 BoNTax validation datasets and eight reference controls. Heatmap of average nucleotide identity values between the validation datasets and the curated *Clostridium botulinum* group I and *Clostridium sporogenes* reference genomes. Higher ANI values are shown in red and lower values in blue. Datasets assigned to *C. botulinum* group I or *C. sporogenes* cluster with their corresponding reference groups, whereas genomes belonging to other *Clostridium* taxa are positioned outside the two target-species groups. The dendrograms represent similarity-based clustering of ANI profiles and not formal phylogenetic relationships.

