## Supplementary figures and images for "BoNTax: a curated genome-based framework for taxonomic classification and characterization of *Clostridium botulinum* Group I and *Clostridium sporogenes*"

### Supplementary Figure 1

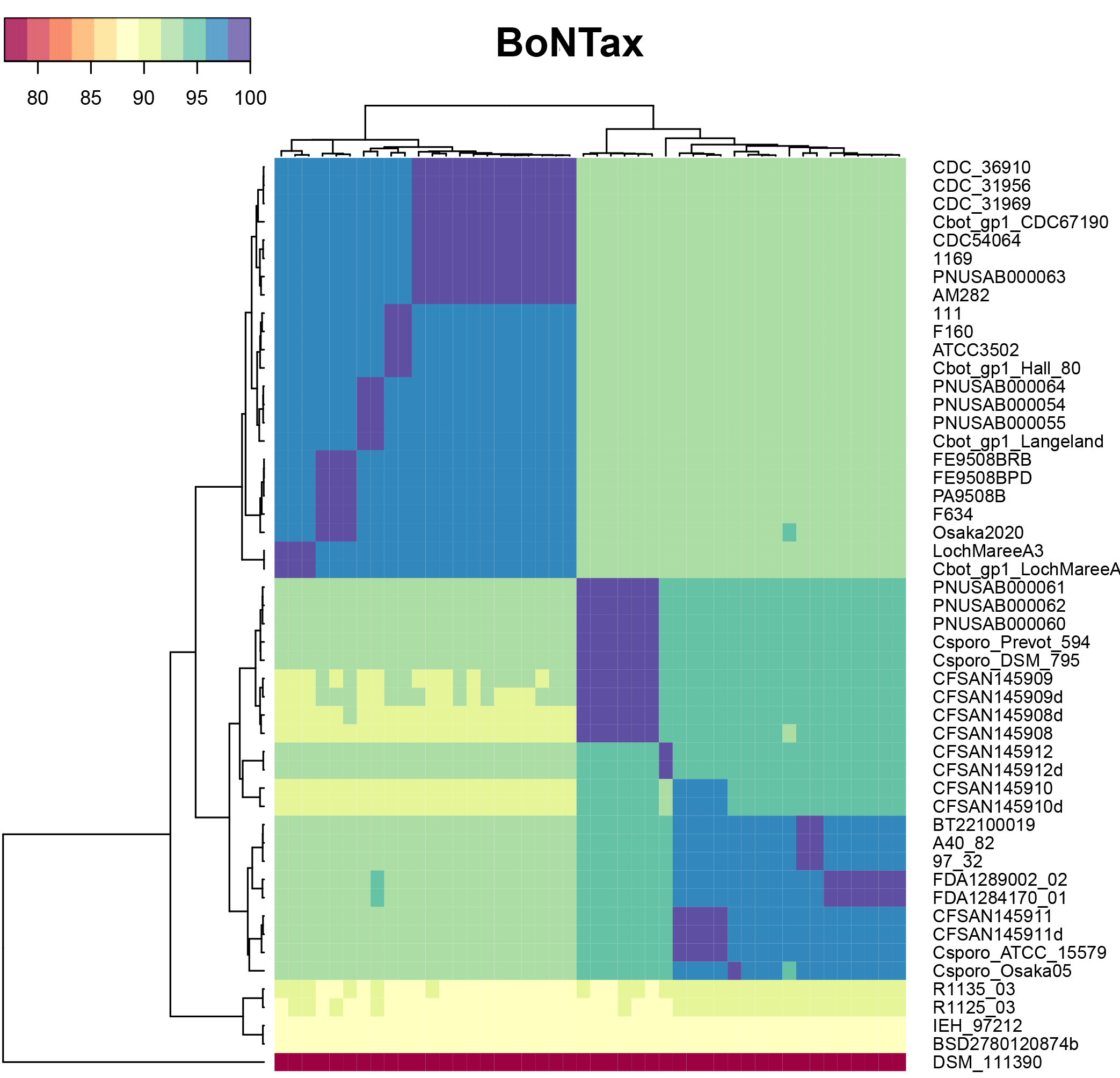
